# The Impact of Low Protein Diet on the Molecular and Cellular Development of the Fetal Kidney

**DOI:** 10.1101/2023.12.04.569988

**Authors:** Kieran M. Short, Giovane G. Tortelote, Lynelle K. Jones, Fabiola Diniz, Francesca Edgington-Giordano, Luise A. Cullen-McEwen, Jan Schröder, Ashley Spencer, Andrew Keniry, Jose M. Polo, John F. Bertram, Marnie E. Blewitt, Ian M. Smyth, Samir S. El-Dahr

## Abstract

**Background:** Low nephron number has a direct impact on the development of hypertension and chronic kidney disease later in life. While intrauterine growth restriction caused by maternal low protein diet (LPD) is thought to be a significant cause of reduced nephron endowment in impoverished communities, its influence on the cellular and molecular processes which drive nephron formation are poorly understood.

**Methods:** We conducted a comprehensive characterization of the impact of LPD on kidney development using tomographic and confocal imaging to quantify changes in branching morphogenesis and the cellular and morphological features of nephrogenic niches across development. These analyses were paired with single-cell RNA sequencing to dissect the transcriptional changes that LPD imposes during renal development to affect nephron number.

**Results:** Single cell analysis at E14.5 and P0 revealed differences in the expression of genes and pathways involved in metabolism, cell cycle, epigenetic regulators and reciprocal inductive signals in most cell types analyzed, yielding imbalances and shifts in cellular energy production and cellular trajectories. In the nephron progenitor cells, LPD impeded cellular commitment and differentiation towards pre-tubular and renal vesicle structures. Confocal microscopy revealed a reduction in the number of pre-tubular aggregates and proliferation in nephron progenitor cells. We also found changes in branching morphogenesis, with a reduction in cell proliferation in the ureteric tips as well as reduced tip and tip parent lengths by optical projection tomography which causes patterning defects.

**Conclusions:** This unique profiling demonstrates how a fetal programming defect leads to low nephron endowment which is intricately linked to changes in both branching morphogenesis and the commitment of nephron progenitor cells. The commitment of progenitor cells is pivotal for nephron formation and is significantly influenced by nutritional factors, with a low protein diet driving alterations in this program which directly results in a reduced nephron endowment.

**Significance Statement:** While a mother’s diet can negatively impact the number of nephrons in the kidneys of her offspring, the root cellular and molecular drivers of these deficits have not been rigorously explored. In this study we use advanced imaging and gene expression analysis in mouse models to define how a maternal low protein diet, analogous to that of impoverished communities, results in reduced nephron endowment. We find that low protein diet has pleiotropic effects on metabolism and the normal developmental programs of gene expression. These profoundly impact the process of branching morphogenesis necessary to establish niches for nephron generation and change cell behaviors which regulate how and when nephron progenitor cells commit to differentiation.

## Introduction

Nephron endowment has a direct impact on renal function and is inversely related to the risk of hypertension and adult kidney disease ^1–3^. Endowment can vary 13-fold in normal human populations ^4^ but the intrinsic drivers of these differences are not fully known. While genetic factors no doubt account for some of this variation (different mouse strains have different endowment ^5^), maternal disease and diet during pregnancy are also likely to significantly influence this measure as all nephrons in humans (and most in mice) are formed *in utero*. One such dietary variable is protein deficiency, which has been identified as a common global nutritional deficit ^6–8^ and one that has been exacerbated by the recent COVID-19 pandemic ^9^. However, the precise molecular mechanism by which maternal low protein diet (LPD) affects embryo and kidney development remains unclear.

Studies in rodent models have shown that an iso-caloric LPD leads to impaired kidney development and reduced nephron endowment (particularly in individuals with low birth weight ^10, 11^) and that it predisposes to the development of hypertension and kidney disease in adult animals ^12–17^. These observations indicate that appropriate quantities of high-quality protein during pregnancy are essential to offset imbalances causative of hypertension and chronic kidney disease later in life ^15, 16^. Therefore, a mechanistic understanding of how genetic and epigenetic changes influence nephron endowment in this context is of clinical importance.

Recent dietary intervention studies have investigated the effects of LPD in rats ^18^ and calorie-restricted diets (CRD) in mice ^19^. These studies report significant reductions in nephron number at birth and nephron progenitor cell proliferation at E14.5. The rat LPD study focused on miRNA targets from whole kidney extracts without addressing the cell specific transcriptional or morphological impact on ureteric tree growth, patterning, and nephron progenitor cell differentiation. Although a different dietary intervention, the CRD study identified methionine metabolism and MTOR signaling in isolated nephron progenitor cells (NPCs) as key drivers of nephron deficiency and showed that a Tsc1 genetic model provided resistance to the CRD kidney phenotype. A limitation of these studies is that they did not address dietary effects on nephrogenesis, ureteric bud involvement and branching, or the cellular transcriptional changes underlying nephron deficiency.

In this study, we report the cellular and molecular mechanisms by which a gestational low-protein diet (LPD) leads to reduced nephron endowment over developmental time. We found that LPD reduces nephron numbers by decreasing cell proliferation early in development and altering nephrogenic and branching morphogenesis programs. This results in fewer nascent nephron structures, such as pretubular aggregates and renal vesicles. Single cell sequencing of LPD kidneys revealed disruptions in cell metabolism and transcription programs related to branching morphogenesis and nephrogenic differentiation. Overall, this study advances our understanding of the cellular and morphological disruptions that contribute to maternally influenced nephron deficits.

## Results

### Maternal LPD reduces postnatal body mass and nephron number

To investigate the impact of LPD on fetal kidney development, male and female mice were fed modified chow for two weeks before conception, throughout pregnancy and until weaning. At embryonic day (E)14.5, body mass was similar between diets and placental mass was reduced by ∼7% (Supplemental Figure S1 and Supplemental Dataset 1). At birth and postnatal day (P)21 body mass in LPD offspring was also lower than normal protein controls (9.5% and 16% respectively, Figure 1A-C). Both newborn (Figure 1D) and 3-week-old LPD mice (Figure 1E) exhibited 19% and 23% fewer nephrons respectively (Supplemental Dataset 1) confirming that the protein-deficient (but isocaloric) diet used in this study causes persistent low nephron endowment and intra-uterine growth restriction. No sex difference in body mass or nephron number were observed.

**Figure 1.**
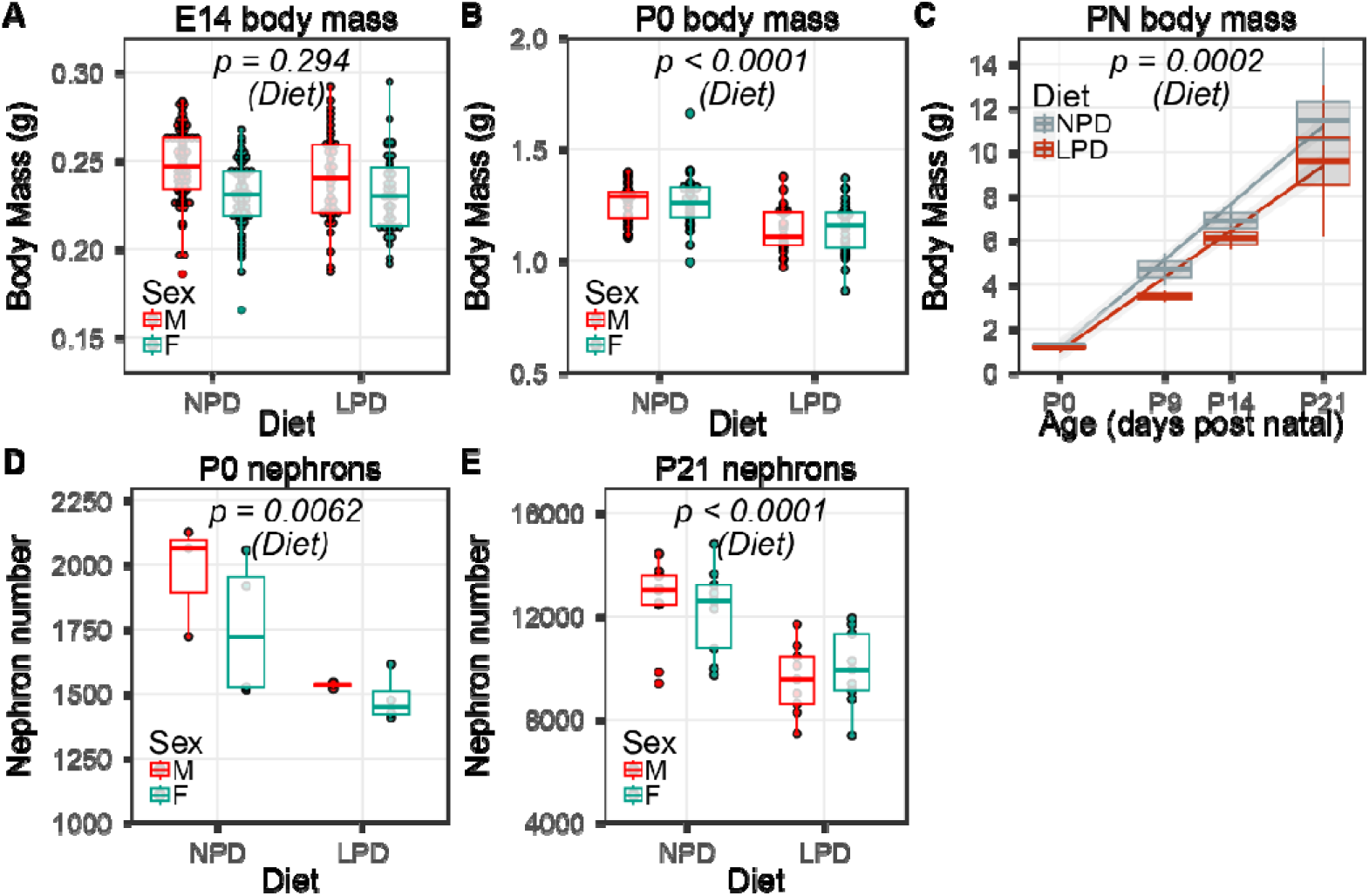
Maternal low protein diet affects the mouse embryo and kidney. At embryonic day E14, body mass is no different depending on diet (A, n=133/86 NPD/LPD, *p = 0.294*), but this changes by birth (B, n = 66/56, *p < 0.0001*). Postnatally, a low protein diet continues to impact postnatal body mass through to PN21 (C, n=133/86 P0, n= 18/15 P9, n= 24/14 P14, n=59/41 P21, NPD/LPD, *p = 0.0002*). At birth, total nephron number is 19% lower in LPD than NPD (D, n=7/7, *p = 0.0062*) and by postnatal day 21, nephron number is 23% lower in LPD (E, n = 18/18, *p < 0.0001*). No sex differences in these parameters were found. Orders for sample numbers are NPD/LPD.

### Maternal LPD alters embryonic kidney branching morphogenesis and nephrogenic niche number

Given the importance of the reciprocal inductive signals between ureteric bud (UB) and nephron progenitor cells (NPCs) during kidney development, we wondered whether branching morphogenesis was affected by LPD and whether such changes contribute to reduced nephron endowment. Branching morphogenesis was assessed at E14.5 using Optical Projection Tomography ^20^ and at P0 by the number of peripheral SIX2+ nephrogenic niches (as a measure of the extent of branching ^21^). While a small 9% difference in niche number was identified at P0 (Figure 2A), at E14.5 the tip number was invariant (Figure 2B, Supplemental Dataset 1). However, the same LPD embryonic kidneys had 25% less volume (Figure 2C) and an 18.4% reduction in surface area (Figure 2D). This leads to an increase in ureteric tip density of approximately 16% in the E14.5 LPD kidneys (Figure 2E). Notably, the terminal branch generations in the LPD offspring were significantly shorter, with the tips reduced by 22.2% and their “parents” by 15% (Figure 2F). To determine if these changes altered the morphology of the ureteric tree, we compared our data with previously defined models of kidney branch patterning ^22^. Branching is a dynamic process and neighboring end branches and their tips normally assume one of three local patterns, based on the how the tips are inherited from parental branches ^22^. In normal kidneys, most branching exhibits a mild asymmetry (“Half Delay” pattern) whereas other more asymmetrical (“Fibonacci”) or perfectly symmetrical (“Perfect”, as in a simple bifurcation) patterns are less common. However, in the LPD kidneys, there are a greater number of imbalanced ‘Fibonacci’ patterned terminal branches (Figure 2F). Similar increases have been noted in kidneys with increased tip packing at the organ periphery ^22^. We therefore assessed nearest-neighbor distances between tips and found this to be ∼25% less in LPD offspring (Figure 2G). This change is analogous to that observed in *Spry1* mutant mice which also have decreased organ size but no change in branching ^22^. These defects in ureteric branching, growth, patterning and tip packing may influence (or be influenced by) nephron formation.

**Figure 2.**
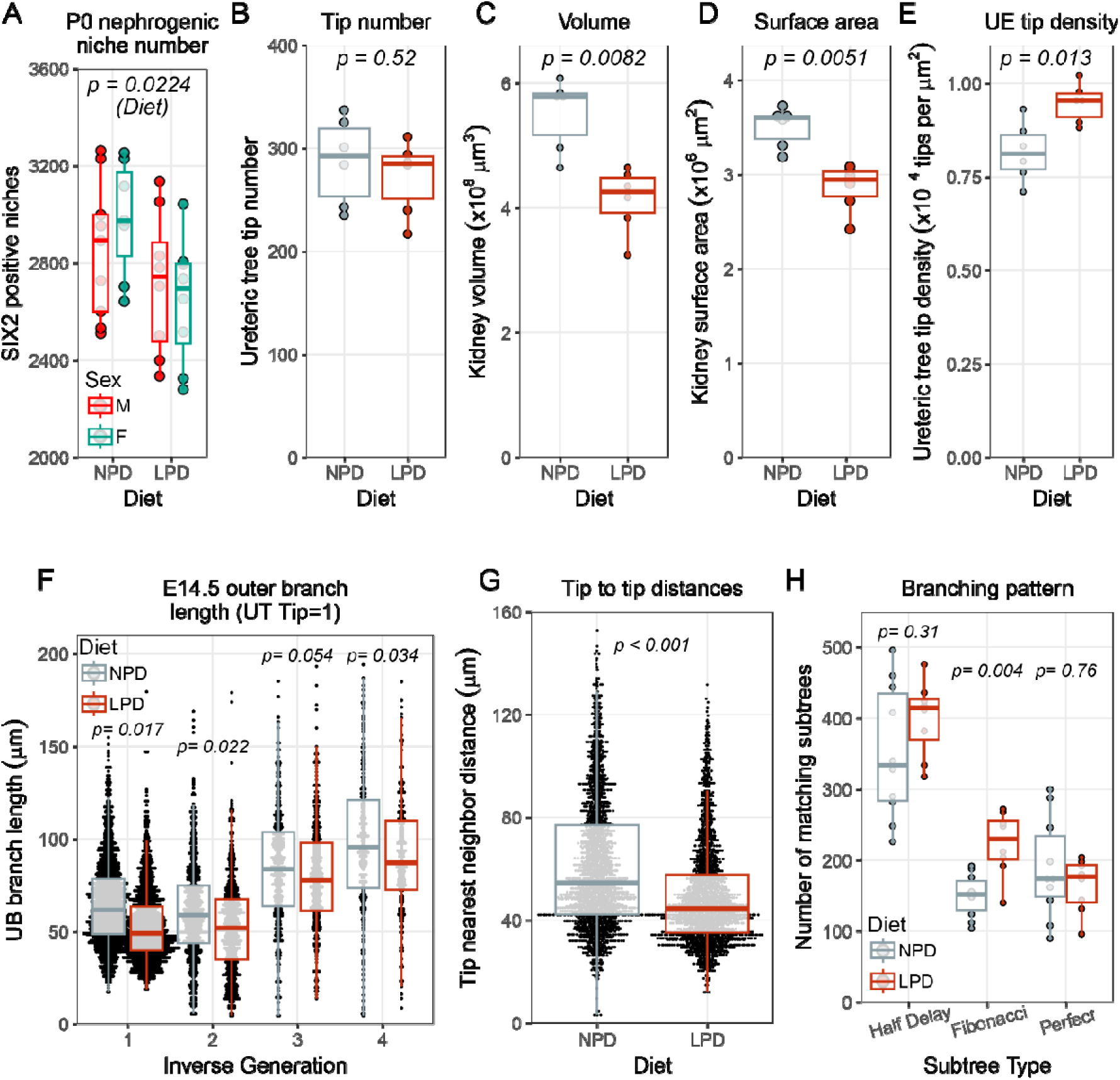
The impact of LPD on branching morphogenesis and kidney growth. At the day of birth, the number of Six2 positive nephrogenic niches is lower in LPD pups and there are no differences between the sexes (A, n=16/16, *p = 0.0224* for diet). This represents a reduction in tip number of the ureteric tree. Although earlier at embryonic day E14.5, the tip number is unchanged (B, *p = 0.52*) but those tips are packed into a smaller kidney (C, *p = 0.0082*) with a smaller surface area (D, *p = 0.0051*) leading to a higher tip density (E, *p = 0.013*). The outer branches of the ureteric tree starting at the tips towards the ureter are consistently shorter in length in the LPD kidneys (F, tip *p = 0.012*, tip parent *p = 0.022*, tip great grandparent *p = 0.034*). The tip grandparent is *n.s*, The shorter branches which are packed more densely have shorter tip to tip distances (G, *p = 0.00072*) with less space to grow. This tighter packing partially affects the LPD branching pattern, with the branch patterns at the periphery fitting a Fibonacci tip state model ^22^ at a greater proportion than the NPD kidneys (H, *p = 0.0044*). N = 6/6 NPD/LPD for whole kidney measurements, and branches analyzed per kidney are n > 1000.

### Maternal LPD reduces progenitor cell proliferation and commitment to nephron formation

The differences in branching and kidney size observed in LPD embryos prompted us to investigate whether proliferation of the ureteric and nephron progenitor cells was affected. A deficit in proliferative capacity would have a negative impact on branching and nephron differentiation. We imaged and counted markers for nephron progenitor cells (SIX2), ureteric bud (TROP2), and cell proliferation (PHH3) in wholemount-stained kidneys. Following LPD exposure, nephrogenic niches at E14.5 exhibited a reduction in proliferation in both the ureteric tip epithelium (25.57%) and cap mesenchyme (27.44%) when normalized to the number of cells in the respective compartments (Figure 3A-C). At P0, proliferation in the LPD cap mesenchyme was further reduced (37.5%) but UB proliferation had normalized (Figure 3A-B, Supplemental Dataset 1). By P2, no differences in cell proliferation in either niche were noted (Figure 3A-B). To determine the impact of LPD on nephrogenesis, we quantified the number of developing nascent nephrons at E14.5, P0 and P2 scoring for pretubular aggregates (PTA), renal vesicles (RV), comma-shaped bodies (CSB), S-shaped bodies (SSB) and Stage IV (nearly mature) nephrons. Despite the preservation of nephrogenic tip niches at E14.5, LPD caused a consistent reduction in PTA number (60.2%, 37.3% and 41% respectively) and the total number of nephrons at any stage of differentiation (31.9%, 18.8% and 27.3% respectively) (Figure 3C-D). The number of distally connected nephric segments indicated no differences *in utero*, but by P2 there was a 16.7% reduction in these structures (Figure 3E). These results suggested that LPD affects the ability of NPC cells to commit to nephron formation.

**Figure 3.**
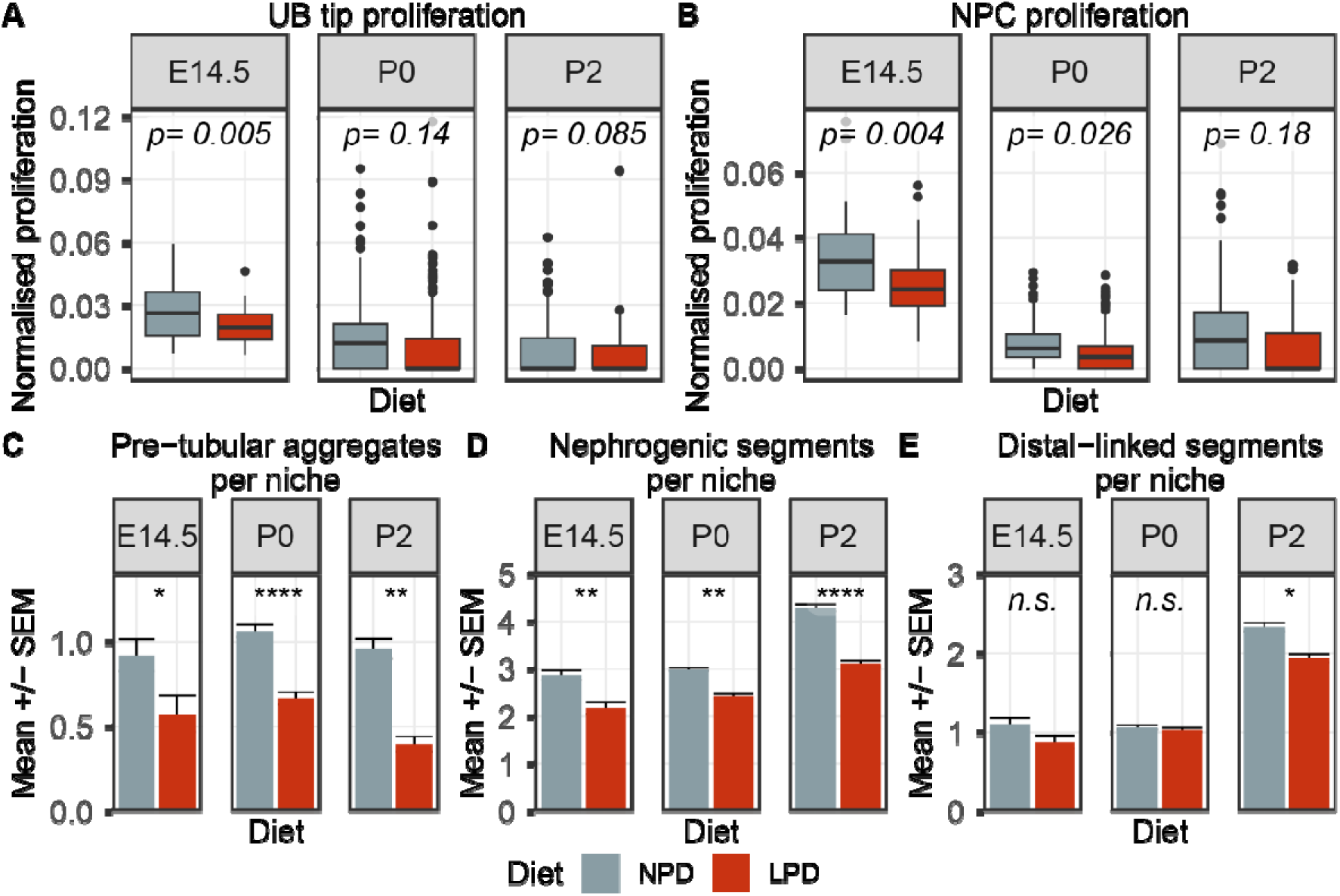
Analysis of proliferation and early nascent nephrons at E14.5, P0 and P2. Proliferation of the ureteric bud tip in LPD offspring is reduced at E14.5 but is indifferent later in development (A), while NPC proliferation is impacted at E14.5 and P0 but not P2 towards the end of nephrogenesis (B). The number of pretubular aggregates (C) and total nephrogenic segments per niche are reduced throughout development in LPD offspring (D). While the number of distal-linked segments per niche appears to be indifferent at E14.5 and P0, they are reduced by P2 (E). Sample numbers, n = 38/40 niches (n = 4/4 kidneys) (NPD/LPD) at E14.5, n = 282/282 niches (9/11 kidneys) (NPD/LPD) at P0, n = 145/144 niches (n=7/7 kidneys) (NPD/LPD) at P2. *p < 0.05, **p < 0.01, ***p < 0.001, ****p < 0.0001, n.s. = not significant.

### Single-cell sequencing reveals developmental and morphological effects of maternal LPD

To better understand the molecular mechanisms underlying these phenotypes, we assessed E14.5 and P0 kidneys by single-cell and single-nuclear RNA sequencing. Quality controlled, filtered and clustered datasets were assigned to different cell types *a priori* using known markers (Figure 4A-D). Diet did not induce an overt change in clustering cell content or UMAP structure (Supplemental Figure S2) and as expected, the P0 kidneys showed more differentiated cellular identities and lower proportions of undifferentiated cell types compared to the E14.5 kidney (Figure 4D). A large proportion of the most variably expressed genes relate to age and are therefore common between the two diets (2968 and 2546 genes at E14.5 and P0 respectively), however dietary treatment altered 1032 and 1454 genes at these timepoints. Preliminary characterization of differential gene expression using gene ontology analysis revealed most changes in these clusters are related to developmental, morphological and metabolic processes (Figure 4F and Supplemental Table S1 and S2) and revealed that the nephrogenic and the ureteric bud lineages were those most impacted by LPD.

**Figure 4.**
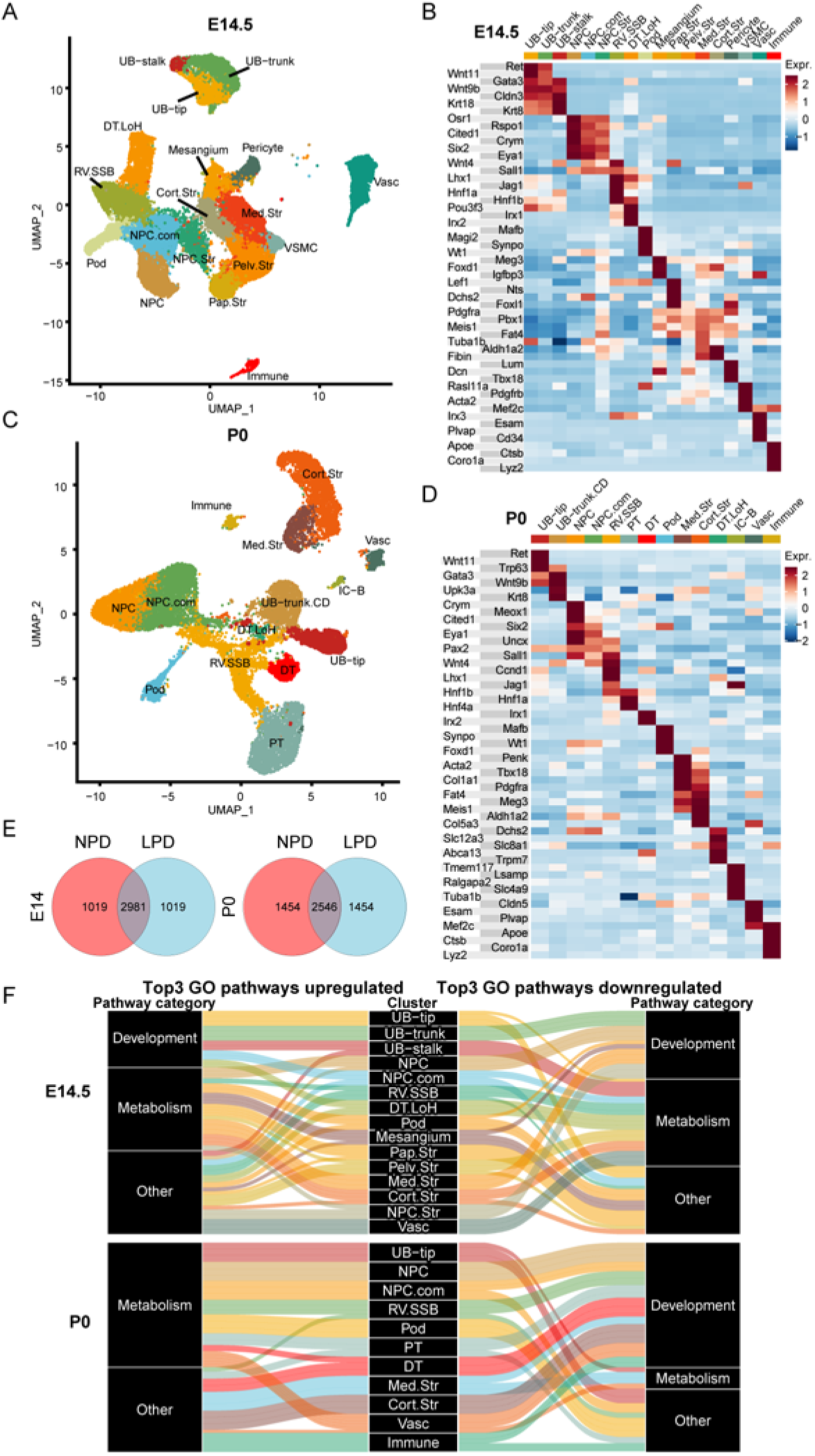
Single-cell and nuclear analysis reveals overarching developmental and metabolic changes in gene expression. Viewed in UMAP reduced dimension space, clusters identified at E14.5 from single-cell sequencing of 30,845 cells (A) partition into known kidney cell types typified by canonical cell type markers (B), and likewise from P0 single nuclear sequencing of 17954 cells (C), the clusters represent known developmental cell types by canonical marker expression (D). Analyzing the most variable genes between NPD and LPD diets at both ages (E) a significant proportion of the most variable genes are unique to the diets. (F) The upregulated (to the left side) and downregulated (to the right side) top GO pathway categories for E14.5 and P0 clusters. Based on the GO term the categories were Development, Metabolism, and everything else, on the right). A significant proportion of the top three GO terms based on the differentially expressed genes in every cluster metabolic effect on the kidney.

### LPD causes disrupted metabolism and epigenetic gene expression profile throughout the kidney

Unifying the impact of the dietary intervention on the cellular metabolism of the developing kidney, we initially saw a consistent disruption in several metabolic pathways in all cells and both ages in response to LPD. While some of the metabolic genes that are differentially expressed in LPD differ between the two ages and cell types within the same kidney, many genes were associated with the same or similar pathways as defined by intersecting GO ontology terms (see Supplemental Dataset 2). The pathways broadly included ATP production, RNA metabolism, and metabolic stress. The genes associated with ATP metabolism included *Atp1a1*, *Atp5c1*, *Cpt1a*, *Gapdh* and *Trp53*. *Trp53* (*p53*) is downregulated in all 18 clusters at E14.5 and plays an important role in the NPC renewal and commitment process ^23^. Upon genetic deletion of *Trp53* ^23^, the most transcriptionally affected gene in NPC cells is *Pck1*. *Pck1*, a dietary responsive gluconeogenesis gene ^24^ which we found was downregulated in the nephrogenic lineages at both E14.5 and P0 (Supplemental Dataset 2).

RNA metabolic processes were also affected by LPD, including genes associated with the regulation of amino acid levels, transcription of ribosomal proteins and translation. Prominent examples included the downregulation of *Hnrnpa0* and *Ybx3* which are involved in the regulation of RNA and amino acid levels (Table I and Supplemental Dataset 2). Alterations in the transcription of ribosomal protein-coding genes (including many of the *Rps/l* family genes) were also detected at both ages, which are indicative of a general impact on transcriptional and translational processes caused by LPD.

Metabolic stress gene expression changes in the embryonic and postnatal kidneys induced by LPD at E14 and P0 as shown in Table I and Supplemental Table S1. We found upregulation of *Glo1*, *Ldhb*, *Dera* and *Rdh10* as well as AP-1 factors like *Fosb* and *Jun* and the transcription factor *Atf4*. ATF4 is known for its role in regulating the expression of genes involved in oxidative stress, amino acid (and protein) synthesis, and cell differentiation ^25^ which all may contribute to the proliferation and differentiation deficits observed in the LPD kidneys.

Epigenetic gene expression exhibited a noteworthy impact, with the most prominent feature being the downregulation of *Ezh2* in LPD kidneys over pseudotime (this analysis discussed later). Specifically, this downregulation was evident in the P0 nephron progenitors, committing nephron progenitors, and across the entire nephron progenitor to podocyte lineage (Supplemental Figure S3, Supplemental Data 1). At E14.5, a significant reduction in *Ezh2* expression was detected across the nephron progenitor to podocyte pseudotime lineage, indicating a potential impact on nephron progenitor maintenance and differentiation (Supplemental Figure S3, Supplemental Data 1). This consistent downregulation of *Ezh2* suggests that LPD may impair the epigenetic regulation necessary for proper nephron progenitor development and transition to podocytes, potentially contributing to the observed phenotypic changes.

### Maternal LPD impairs the differentiation of nephron progenitor cells and alters the expression of developmental and metabolic genes

Using traditional cluster gene expression analysis, we observed downregulation of several genes related to the cell cycle and proliferation in LPD NPCs at E14.5 (*Ybx3, Ptn, Gpc3, Rcc2*) (Table 1), aligning with decreased NPC proliferation (Figure 3B). GO analysis identified terms linked to cell proliferation in metanephric development (GO:0072203; FDR 0.008), suggesting LPD impacts progenitor pool maintenance and development (see Supplemental Dataset 2 and Table S2). In the P0 NPC cluster, LPD downregulated genes such as *Dach1* and *Pdgfc* (Table 1). Dach1 interacts with Eya1 in nascent nephrons, and Pdgfc is expressed in nephron progenitors contacting the ureteric tip at E11.5. Eya1 and Pdgfc are critical for NPC establishment, maintenance, and differentiation. This data suggests that LPD disrupts reciprocal inductive signals during kidney development, possibly causing the observed branching phenotype (Figure 2F). In committing nephron progenitors, genes involved in RNA metabolism, splicing, nuclear division, nuclear chromosome segregation, and mitosis (*Rcc2*, *Cdk2ap1*, *Ccnd1*, and *Hnrnpa*) were differentially expressed at E14.5, indicating ongoing metabolic effects in these cells after transitioning from a naïve NPC state. GO terms related to kidney development and tubule morphogenesis were also identified, with mis-expression of genes such as *Crym* (Table 2 and Supplemental Dataset 2). The upregulation of *Crym* just before epithelization (Table 2) suggests a compromised transition to pretubular aggregate cells and epithelialized renal vesicles. At P0, several genes associated with nephron progenitor identity, including *Pax2*, *Bmper*, *Wt1*, and *Magi2* were downregulated in LPD kidneys (Table 2), supporting the observed reduction in PTAs (Figure 3C).

**Table 1.** E14.5 and P0 DEG between NPD and LPD NPC. The table presents the DEG between NPC from NPD and LPD cohort (cutoff log FC greater than |0.5| and p-value < 0.05). DEG list was generated with EdgeR as described in Methods. p-value threshold for the differences was set as p<0.05. logFC = log of the fold change difference; logCPM= log of counts per million reads mapped; PValue = calculated p-value for differences.

| Gene | LogFC | LogCPM | PValue | Age | Direction |
| --- | --- | --- | --- | --- | --- |
| Hnrnpa0 | -1.318 | 9.277 | 1.49E-84 | E14 | Down |
| **Ybx3** | **-1.090** | **9.351** | **8.79E-88** | **E14** | **Down** |
| Ptn | -0.884 | 9.201 | 6.82E-35 | E14 | Down |
| **Rcc2** | **-0.548** | **9.109** | **2.50E-15** | **E14** | **Down** |
| Ubb | -0.540 | 10.748 | 1.06E-61 | E14 | Down |
| **Gpc3** | **-0.525** | **9.417** | **7.92E-19** | **E14** | **Down** |
| Snhg12 | 0.552 | 9.060 | 5.38E-14 | E14 | Up |
| **Tgfb1** | **0.580** | **9.102** | **1.23E-15** | **E14** | **Up** |
| Erh | 0.589 | 9.130 | 1.67E-17 | E14 | Up |
| **Crym** | **0.990** | **9.655** | **3.99E-65** | **E14** | **Up** |
| Glo1 | 1.178 | 9.389 | 1.12E-76 | E14 | Up |
| **Aldh1a2** | **1.284** | **9.000** | **7.46E-53** | **E14** | **Up** |
| Rbfox1 | -0.950 | 12.154 | 2.14E-55 | **P0** | Down |
| **Lhfp** | **-0.840** | **11.150** | **3.04E-23** | **P0** | **Down** |
| Cdh4 | -0.821 | 11.245 | 3.39E-22 | **P0** | Down |
| **Dach1** | **-0.769** | **11.188** | **1.39E-21** | **P0** | **Down** |
| Pdgfc | -0.744 | 11.482 | 7.81E-25 | **P0** | Down |
| **Mdga2** | **-0.670** | **11.177** | **5.76E-14** | **P0** | **Down** |
| Strbp | 0.422 | 11.213 | 2.80E-08 | **P0** | Up |
| **App** | **0.427** | **11.269** | **9.03E-09** | **P0** | **Up** |
| Malat1 | 0.435 | 16.608 | 1.72E-47 | **P0** | Up |
| **Fgfr2** | **0.444** | **11.221** | **4.88E-09** | **P0** | **Up** |
| Hdac8 | 0.523 | 11.345 | 5.44E-12 | **P0** | Up |
| **H3f3b** | **0.568** | **11.212** | **1.42E-13** | **P0** | **Up** |

**Table 2.**
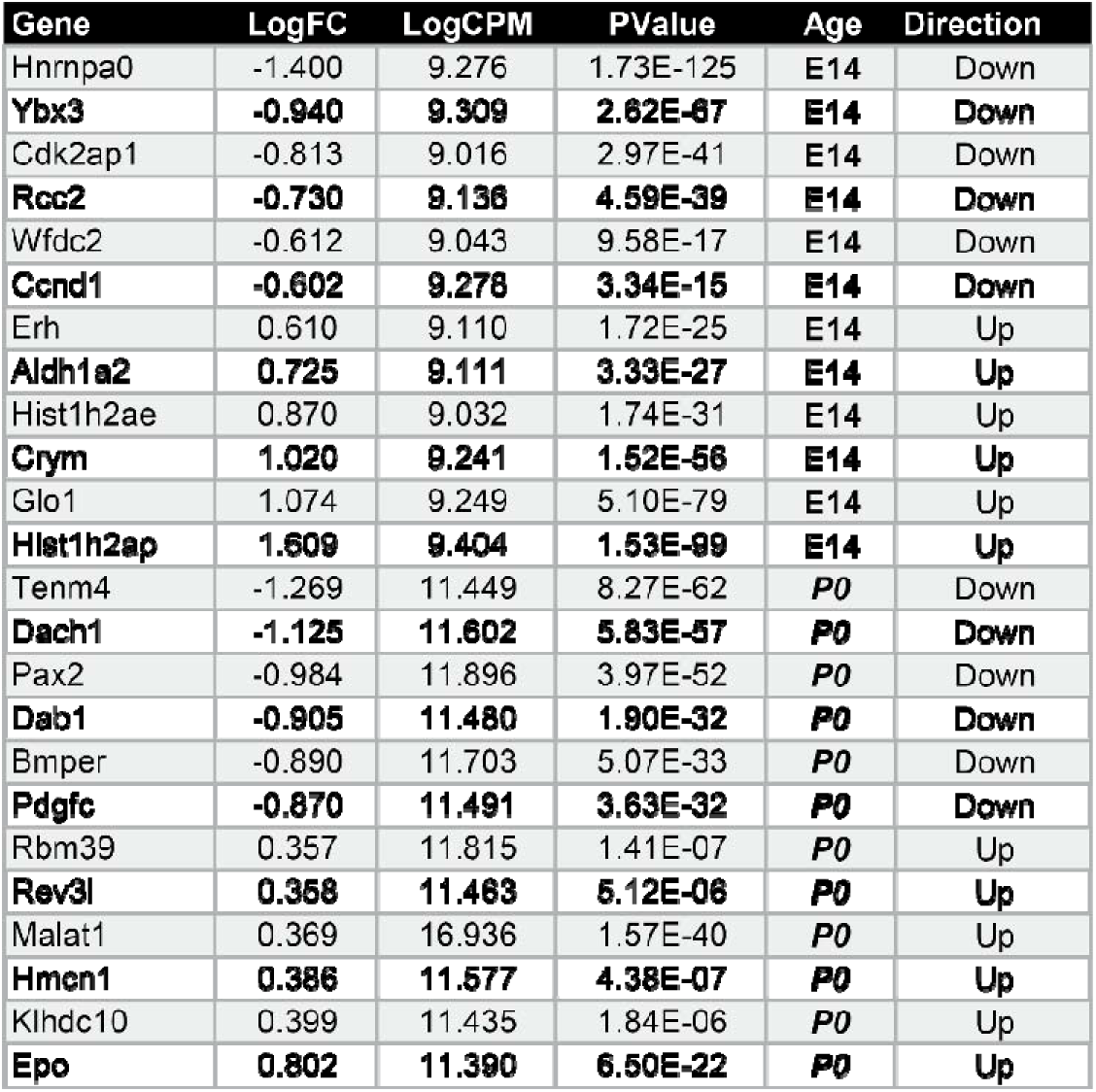
E14.5 and P0 DEG between NPD and LPD NPC comm cluster. The table presents the DEG between NPC comm from NPD and LPD cohort (p-value < 0.05 and cutoff log FC greater than |0.5|). The DEG list was generated with EdgeR as described Methods. p-value threshold for the differences was set as p<0.05. logFC = log of the fold change difference; logCPM= log of counts per million reads mapped; PValue = calculated p-value for differences.

To assess the impact of LPD on the differentiation of NPCs through to podocytes, we next examined cells across the entire nephrogenesis lineage (Figure 5A) before focusing on early NPC differentiation. Pseudotime analysis with Slingshot enabled us to infer an unbiased timeline for all cells in the nephron progenitor lineage without predefined developmental identities. Plotting each cell’s pseudotime predicted value grouped by canonical clusters (Figure 5B) confirmed that this analysis correctly ordered the cells according to the established nephron differentiation program. We then used tradeSeq to investigate dynamic gene expression changes during early nephron formation, clustering genes on a per-cell basis using weighted gene expression correlation (Figure 5C). This analysis revealed downregulated metabolic genes over pseudotime (Figure 5C, E14.5 clusters 8 and 4) and downregulated cell cycle and proliferation genes (Figure 5C, P0 cluster 6) (see Supplemental Dataset 2 for comprehensive gene lists and GO terms). In the context of kidney development, genes crucial for NPC commitment and renal vesicle formation (*Pax2, Fgf8, Wnt4*) exhibited reduced or pseudotime-shifted expression in LPD kidneys in E14.5 and P0 kidneys (Figure 5D, Supplemental Dataset 2). Furthermore, *Notch2* was downregulated in the E14.5 trajectory, and *Jag1* (Figure 5H) and *Notch3* reduced in both E14.5 and P0 trajectories, suggesting a decrease in NOTCH and WNT signaling. This conclusion was supported by decreased WNT4 protein levels in P0 LPD kidneys (Figure 5E). GO ontology analysis revealed an enrichment of Wnt signaling genes (GO terms: 0016055 and 0030177) in clusters 4 and 6 (Figure 5C), supporting the involvement of the generalized Wnt pathway through the NPC to Podocyte trajectory at E14.5 and P0. When analyzing gene expression in the pseudotime of NPC and committing NPC clusters separately, we also identified the ’Wnt signaling pathway’ (GO:0016055) and ’regulation of Wnt signaling pathway’ (GO:0030111, Supplemental Dataset 2) as commonly downregulated across the NPC and committing NPC trajectories at E14.5 and P0. This indicates the potential for a prominent Wnt deficiency caused by the low protein diet.

**Figure 5.**
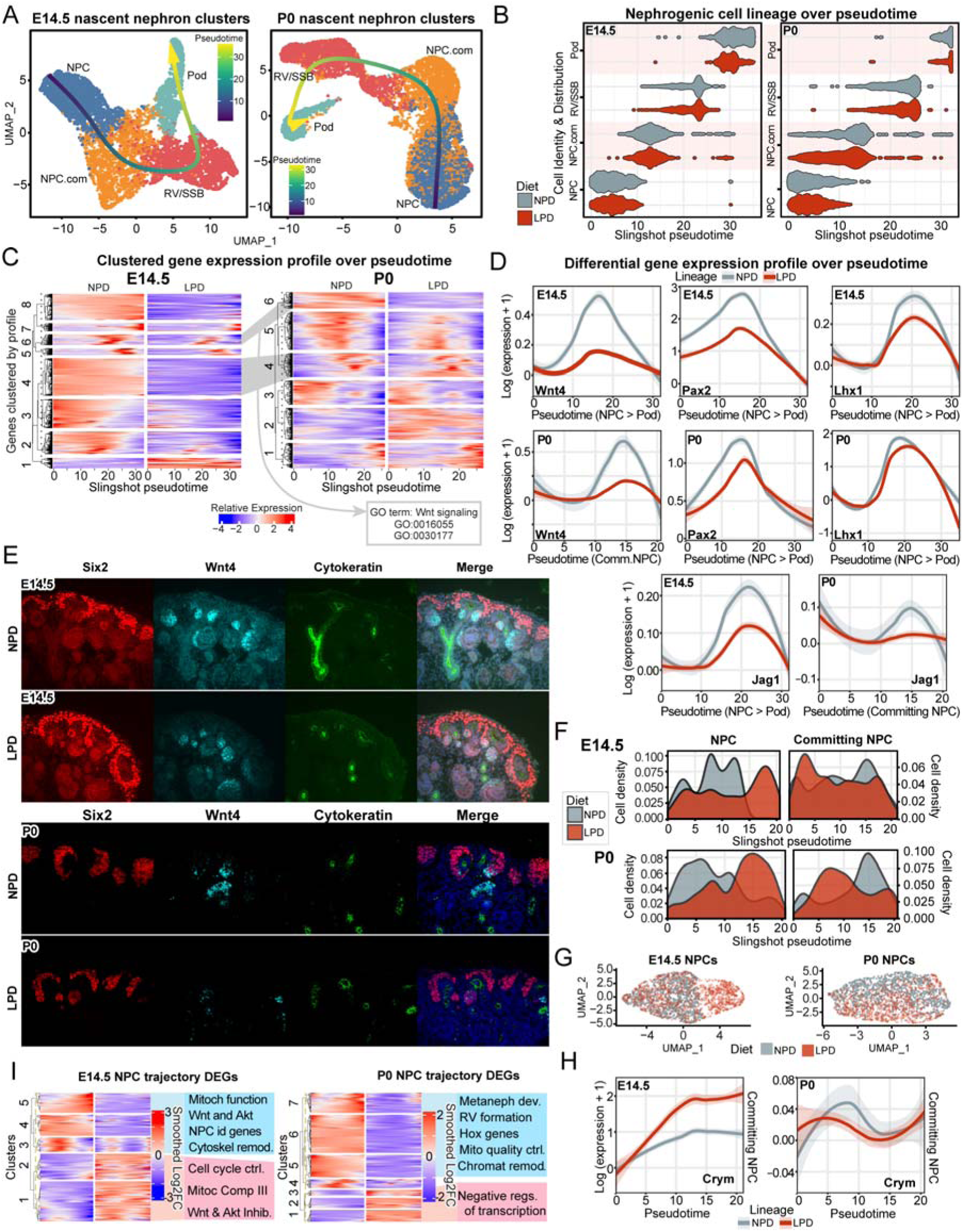
Analysis of the nephrogenic lineage indicates changes in cell development and nephrogenesis-promoting gene expression. NPC, committing NPC (NPC.com), renal vesicle – s-shaped body (RV/SSB) and Podocytes were subset from the whole kidney dataset and reclustered (A). Slingshot was used to infer pseudotime and the resulting timeline recapitulated known nephrogenic progression from NPC to Podocytes (B). Clustering of the dynamic changes in expression of the differentially expressed genes across pseudotime (C) indicates half of the clusters represent genes involved in cellular metabolic and nephrogenesis pathways, including Wnt pathways. *Wnt, Pax2, Lhx1* and *Jag1* genes important for nephrogenesis are downregulated over pseudotime (D). Expression of WNT4 (cyan) in the developing nephrons is reduced in the E14.5 and P0, with nephron progenitor cells marked by SIX2 (red), ureteric tree by cytokeratin (green) and DNA (blue). Cell density plotted over pseudotime reveals an enrichment of LPD cells at the transition between NPC and committing cell identities at E14.5 and P0 (F). Increased expression of Crym at the end of pseudotime indicates the epithelialization of the committing NPCs is reduced (H). Changes in gene expression across the NPC trajectory indicate multiple pathways relatde to cell differentiation, specification and cell cycle and mitochondrial function are affected (I).

UMAP projections of subclustered NPCs revealed areas of diet-specific cell enrichment on the UMAP_1/UMAP_2 projection at P0 and a unique LPD-only pool of cells at E14.5 (Figure 5F). The distribution of these early LPD NPCs and committing NPCs was biased across pseudotime, with a greater proportion of NPCs toward the end (D = 0.443, p < 2.2e-16 at E14.5, and D = 0.386, p < 2.2e-16 at P0, KS test) and a greater proportion of committing NPCs at the beginning (D = 0.136, p = 1.502e-11 at E14, D = 0.265, p < 2.2e-16 at P0, Two-sample KS test) (Figure 5G). The differences in gene expression primarily involved cell commitment genes, such as *Cited1* (-0.28 Log2FC) and Crym (0.99 Log2FC, Tables 1-2, Supplemental Figure S4C-D). The upregulation of *Crym* and lowered expression of *Cited1* in LPD NPCs suggested a more advanced differentiation state, which was further supported by the continued upregulation of *Crym* in committing NPCs (Figure 5H). However, despite these markers indicating advanced differentiation, LPD-specific NPCs continued to express naive NPC markers, with very few cells showing expression of established commitment genes like *Wnt4* or *Itga8* (Supplemental Figure S4D). This contrasting expression pattern is reflected throughout NPC pseudotime. At E14.5, there was a notable downregulation of genes within NPC fate determination pathways such as WNT, AKT, and HOX (Figure 5I, E14.5 clusters 3-5) and an upregulation of their inhibitors (Figure 5I, clusters 1-2, Supplemental Dataset 2). By P0, LPD kidneys exhibited a reduction in genes key to epithelialization and renal vesicle formation, including *Magi2*, *Pax2*, *Itga8*, *Itga9*, *Pbx1*, and *Shroom3* (Figure 5I, P0 clusters 5-7, Supplemental Dataset 2). These changes in gene expression correlate with morphological alterations such as decreased cell proliferation (Figure 3B) and reduced differentiation in the progenitor niche, indicating that LPD affects both nephron progenitors and committing nephron progenitors over pseudotime. The disruption in gene regulation essential for the transition from NPC to committed NPC at both developmental stages likely underlies the observed accumulation of LPD cells at these critical transition stages.

### The branching and proliferation phenotype in the ureteric epithelium have underlying deficits in proliferative and developmental pathways

Given the restricted ureteric branch growth morphology and the differences observed in NPC commitment, we focused on the known determinative role of the ureteric epithelium (UE) in the NPC commitment process. We examined gene expression in the branching ureteric bud (UB) as a potential factor in the impediment of NPC commitment (Figure 6A-B). Growth and bifurcation of the ureteric tips is critical for triggering NPC differentiation, and the observed reduction in ureteric tip lengths and cellular proliferation at E14.5 was reflected in the reduction in proliferation and mitosis gene expression at E14.5 (*Hnmpa0*, *Ybx3*, *Rcc2*, *Gpc3*, *Cdk2ap1*, see Table 3). The decreased growth is also associated with metabolic stress with upregulation of *Glo1* and *Aldh1a3* ^26, 27^. Pseudotime analysis correctly ordered cells from UB-tip through to Stalk or trunk/collecting (Figure 6C) and revealed differences in cell distribution (Figure 6C-D) as well as a reduction in *Wnt9b* expression in LPD cells with a mostly trunk/collecting duct identity at P0 (Figure 6E). Wnt9b triggers *Wnt4* expression ^28^ to initiate NPC commitment, and is normally expressed in the base of the ureteric bud and trunk. While immunofluorescent staining did not reveal any major changes in WNT9B levels (Supplemental Figure S6), *Wnt9b* heterozygote mice exhibit a weaker induction of WNT4 ^28^. This aligns with observations in our study where decreased *Wnt4* expression and delayed NPC commitment occur in LPD kidneys. Broader assessment of gene expression in the ureteric clusters and across pseudotime trajectories revealed impacts on metabolic, cell cycle, and other developmental genes (Figure 6E-F, Supplemental Data S2). These differences were associated with cell proliferation (*Hdac9*, *Erbb4*, *Fgf12*) (Table 3), cytoskeleton remodeling and cell adhesion (*Shroom3*, *Tenm2*, *Ctnnd2*, *Robo1*, *Nrg3*) (Supplemental Data S2). Together, these changes indicate an effect on ureteric tip growth and inductive signal expression brought about by disruption in metabolic and cell cycle gene expression. This results in the reduced cellular proliferation observed at E14.5, and reduced expression of Wnt factors necessary for nephron progenitor commitment.

**Figure 6.**
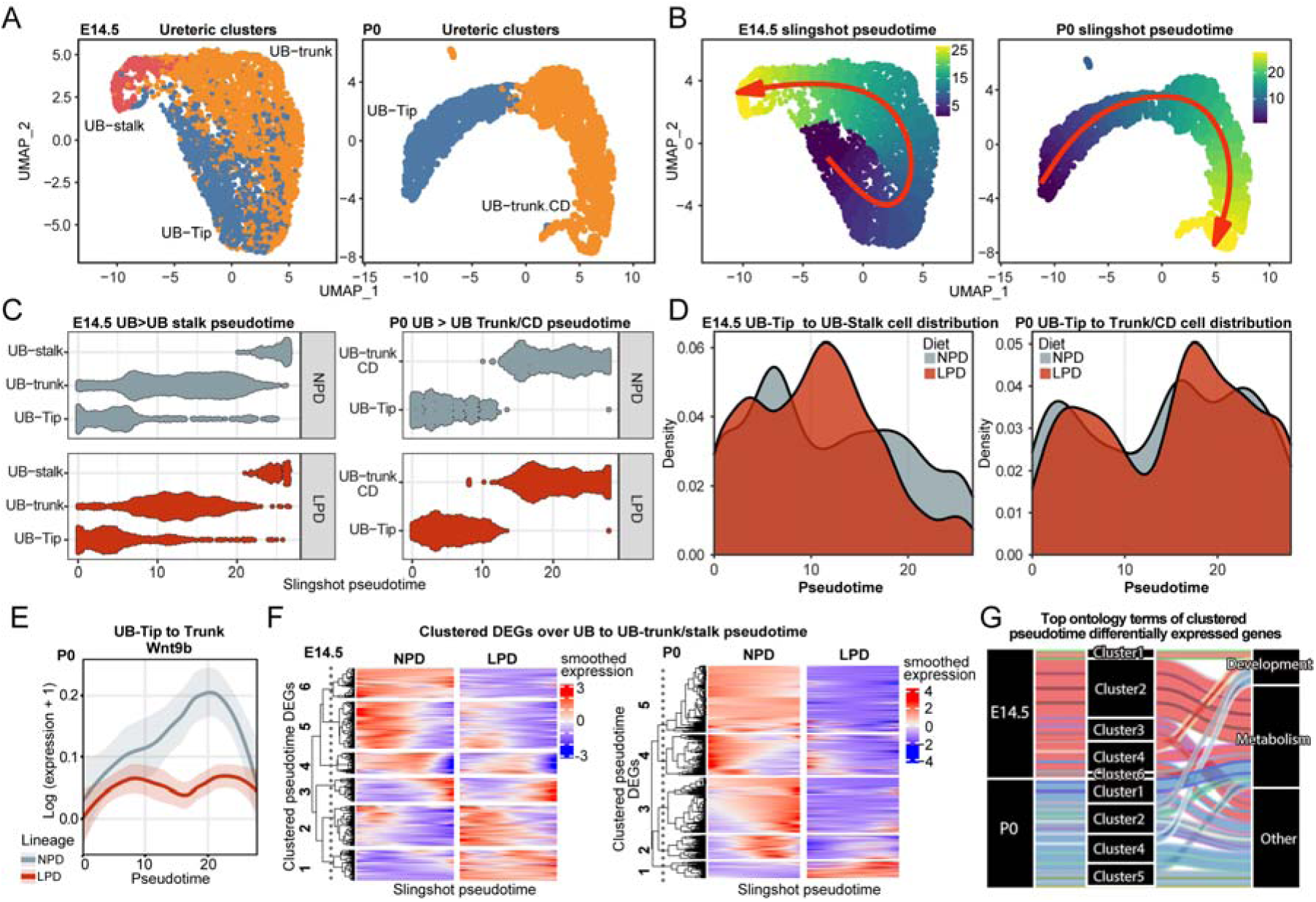
Single-cell analysis of ureteric development at E14.5 and P0. The ureteric lineage from tip to stalk in early development (E14.5) and tip to trunk/collecting duct at P0 was subset from the entire single-cell dataset and re-clustered (A). Pseudotime trajectory analysis followed the established developmental progression of ureteric development (B, C). The distribution of cells indicates a greater proportion of LPD cells in the middle of pseudotime at the approximate point of trunk differentiation (D = 0.14379, p-value < 2.2e-16 at E14.5, and D = 0.086943, p-value = 0.0003057 at P0, KS test) (D). Wnt9b, an NPC commitment and renal vesicle specification gene was downregulated near the middle of pseudotime (E). Analysis of differential gene expression across pseudotime (F) revealed many genes with similar expression profiles having altered expression, with the majority of genes clustered into GO metabolism pathways (G, presented in a manner similar to Figure 4F).

**Table 3.**
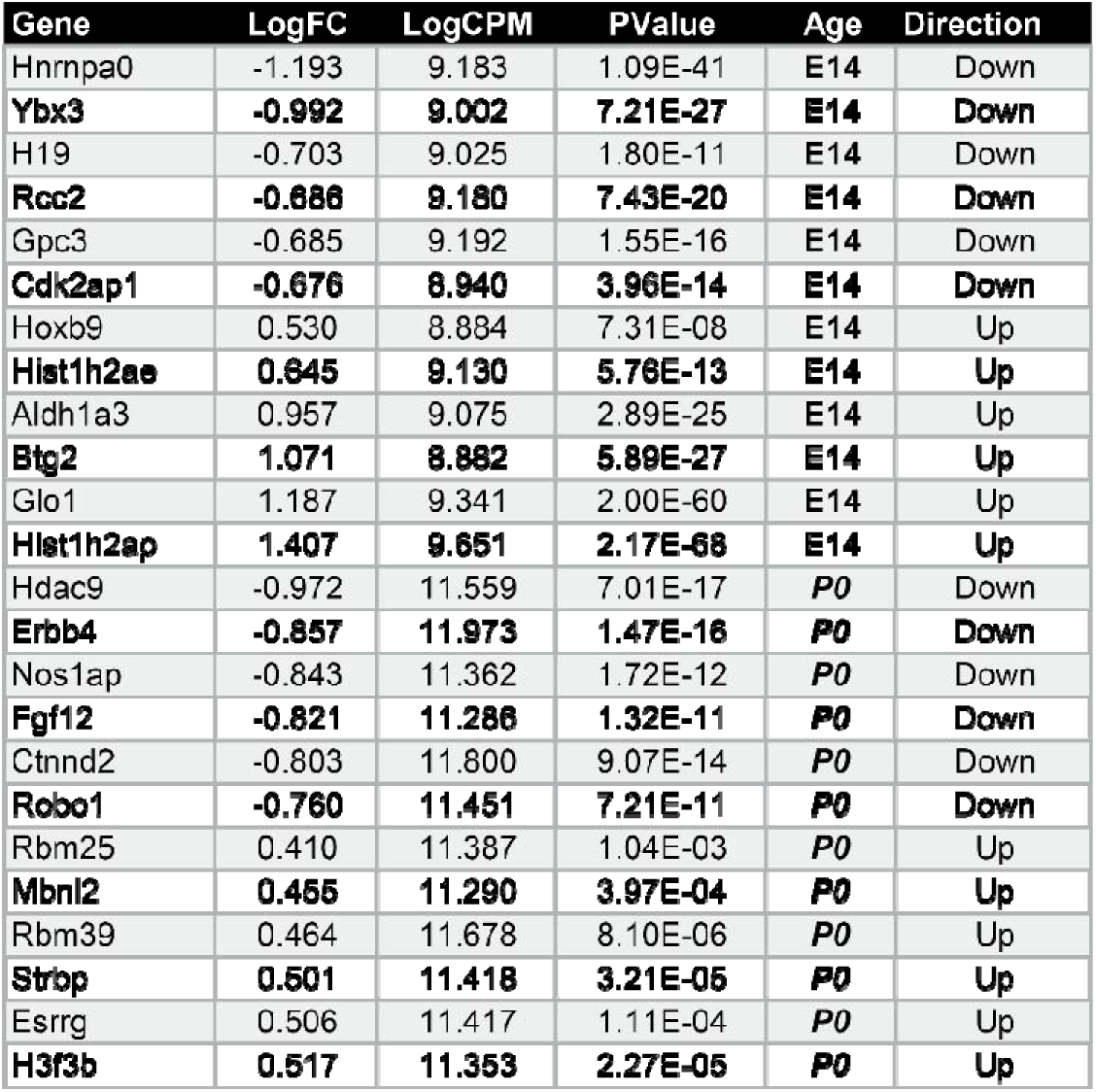
E14.5 and P0 DEG between NPD and LPD UB-tip clusters. The table presents the DEG between UB-tip from NPD and LPD cohort (p-value < 0.05 and cutoff log FC greater than |0.5|). The DEG list was generated with EdgeR as described in Methods. p-value threshold for the differences was set as p<0.05. logFC = log of the fold change difference; logCPM= log of counts per million reads mapped; PValue = calculated p-value for differences.

## Discussion

Nephron deficit is a key feature of intrauterine growth restriction caused by maternal malnutrition. We used a dietary model with reduced protein intake (maintaining caloric balance) to induce a ∼25% reduction in nephron number at birth. This model helped us explore the cellular and molecular bases for reduced nephron endowment. The LPD diet induced significant changes in metabolism gene regulation and cellular metabolism generates the necessary energy, amino acid balance, transcriptional and translational platform for development in all organisms ^29^. In almost every kidney cell type, downregulating metabolic stress, RNA metabolism, and ATP production genes throughout the kidney and in the nephrogenic lineage. This downregulation is crucial as energy production genes play direct roles in NPC commitment ^30^ and nephron number ^31^.

Branching morphogenesis of the ureteric epithelium and maintenance and commitment of nephron progenitor cells are interdependent processes essential for nephron endowment ^28, 32, 33^. Despite reducing kidney size, LPD does not impact the capacity of the ureteric epithelium. Instead, mid-gestation kidneys have shorter branches and higher tip density, leading to patterning defects. Metabolic demands may limit cell proliferation during ureteric epithelial morphogenesis ^34^ and this likely contributes to the UB development abnormalities observed. Consistent with this finding, the shortened ureteric trunk and outer branch generations in LPD kidneys in linked to altered expression of several cell cycle genes. We observed an upregulation of *Bgt2* (a cell cycle repressor ^35^) and reductions in *Shroom3* and *Errb4* in ureteric trunk cells, which are changes known to affect cellular remodeling and duct cell polarization ^36, 37^. Most RET pathway genes controlling UB branching and growth were unaffected, but *Gdnf* and its co-receptor *Gfra1* were downregulated at E14.5, along with branching induction genes (*Dach1* ^38, 39^ and *Ptn* ^40^) expressed elsewhere in the kidney. As epithelial cells transition from the UB-tip to the collecting duct, they transiently express genes central to nephron induction. Pseudotime analysis of UB maturation revealed reduced *Wnt9b* expression as cells progress, which is significant because Wnt9b deletion alters cell movement and polarity within the ureteric trunk ^41^. Wnt9b also acts as a paracrine signal regulating adjacent primed nephron progenitor cells ^42^ and induces Wnt4 expression in committing nephron progenitors, pretubular aggregates, and renal vesicles ^28^. Wnt4 further acts in an autocrine and paracrine manner to sustain its expression ^43^. We found widespread deficiencies in this pathway in LPD kidneys, including reduced *Wnt4* expression in advanced pretubular aggregates/renal vesicles and accumulation of PTAs, whose differentiation depends on normal WNT signaling.

In the nephrogenic niche, LPD significantly reduces progenitor cell commitment to pretubular aggregates, affecting mature nephrogenic structures. Notably, the number of mature “connected” nephrons remains unchanged during embryonic development, suggesting the first round of nephron formation is unaffected by LPD. However, a ∼50% reduction in PTAs during later development correlates with impaired nephron formation at birth, indicating a persistent defect in NPC differentiation. Pseudotime analysis of NPC differentiation revealed significant impacts on WNT and NOTCH pathways, including reduced expression of metanephric development and RV formation genes, which mirror reductions in Wnt9b expression in the developing UB-tip/trunk, affecting renal vesicle genesis. Collectively our analysis suggests the formation of a unique NPC subpopulation with mixed identity following LPD exposure, whose commitment to nephron differentiation is hindered. These cells exhibit increased expression of hallmarks of more differentiated NPCs (*Wnt4*, *Crym*, *Ubx3*) while also co-expressing uncommitted cell marker (like Cited1). This unique cell population clustered with other NPCs, whereas committed LPD and NPD NPCs clustered separately, suggesting they are at the brink of differentiation but fail to commit and transition to PTAs. This is consistent with the deficit in these structures in LPD-exposed embryos. Although a similar unique subset of LPD cells was not evident in P0 NPCs, pseudotime analysis still profiled less differentiated LPD NPCs that gather at the end of NPC pseudotime and the beginning of committing NPC pseudotime, despite reduced proliferation and similar total cell numbers. Guo et al. supports the idea that dietary interventions can block progenitor differentiation, showing how canonical WNT signaling levels regulate the balance between self-renewal and differentiation in NPCs ^44^.

Our study revealed a significant reduction in *Ezh2* gene expression along the NPC lineage trajectory. Ezh2, a crucial epigenetic regulator, is the sole histone lysine 27 methyltransferase in mouse NPCs ^45^. The loss of Ezh2 function in NPCs results in premature progenitor commitment, leading to the expression of genes such as *Wnt4* ^45^, but ultimately fails to drive differentiation into PTAs and RVs. A similar disruption in differentiation into PTAs and RVs is also observed in the LPD nephron lineage. However, in the LPD scenario, *Ezh2* expression is reduced rather than completely ablated, differentiation does not completely fail, and *Wnt4* expression is reduced. The differences in Wnt4 response suggest that the disruption in LPD occurs earlier in the commitment timeline, prior to *Wnt4* expression, whereas *Ezh2* loss may interfere with commitment at a later stage when *Wnt4* is already expressed. Mechanistically, Ezh2, which is abundantly expressed in NPCs, maintains NPC self-renewal through deposition of H3K27me3 and closed chromatin in poised differentiation genes and cell cycle inhibitors such as p16 ^45^.

A recent study employing a low-protein diet (LPD) reported a 28% reduction in nephron numbers at birth in Wistar rats ^18^. The researchers explored the role of microRNAs in nephron deficiency by sequencing miRNAs from embryonic day 17 rat kidneys—comparable to E15 in mice—and subsequently performed immunostaining of miRNA target proteins. In contrast, our single-cell sequencing indicated stable expressions of *mTor*, *Tgfb1* (increased in their study), *cmyc*, and *Ki67* (both decreased in their analysis). Beta-catenin was upregulated in their findings, whereas our results showed minimal and inconsistent changes in gene expression at E14.5. Interestingly, NPC *Zeb1* gene expression increased by 0.22 Log2FC in our study, aligning with a 30% increase in ZEB1 they reported. Their approach of using sections to count NPC cell numbers identified a reduced count in tip/cap niches. However, our comprehensive analysis of every cell in every niche by 3D imaging did not find any differences in cell number and suggests that sectional counts may not capture the full complexity of NPC cell distributions. Similar to our observations in the E14.5 cohort, their study observed reductions in nephron numbers and progenitor cell proliferation in LPD UB-tip and NPCs. However, it did not address several critical aspects covered in our research, such as impacts on UB-tip growth and ureteric tree patterning, normalization of NPC and UB-tip proliferation by P2, and the extensive transcriptional and morphological changes affecting nephron progenitor commitment and differentiation.

Another study utilizing a calorie-restricted diet (CRD) in mice^19^ highlighted the involvement of specific pathways such as methionine metabolism and mTOR signaling through mTorc1 in isolated nephron progenitor cells (NPCs), focusing primarily on molecular and cellular responses in vitro. This research progressed to in vivo interventions with maternal methionine supplementation and genetically enhancing mTorc1 signaling, which recovered nephron endowment. Both maternal CRD and LPD cause a reduction in body mass and postnatal nephron number, with NPC proliferation reduced at E14.5 in both cases. Unlike the CRD study, our study did not detect changes in mTOR signaling or *mTorc1* in low-protein diet (LPD)-treated kidney NPCs at critical developmental stages (E14.5 or P0), suggesting a distinct metabolic response to protein restriction compared to calorie restriction. Furthermore, a key difference emerged in the recovery of NPC proliferation: in CRD-treated kidneys, proliferation normalized by birth, whereas in LPD-treated kidneys, normalization was delayed until postnatal day 2 (P2). This discrepancy may reflect the more severe or prolonged developmental hurdles faced by NPCs under protein stress. Additionally, both studies demonstrated an early reduction in cell cycle activity; however, the quicker normalization in the CRD model points to potential differences in how metabolic and protein deficiencies impact cellular recovery and development.

Our findings demonstrate that LPD reduces nephron numbers through disruptions in cell proliferation, branching morphogenesis, and nephrogenic differentiation, culminating in fewer nascent nephron structures such as pretubular aggregates and renal vesicles. Single-cell sequencing revealed significant changes in cell metabolism and transcription programs, particularly impacting nephrogenic and ureteric bud lineages. These alterations correspond to a decreased expression of key developmental regulatory genes and pathways, notably in Wnt signaling, which are crucial for nephron progenitor commitment and differentiation. The broader implications of this study highlight the long-term utility of these findings in understanding the role of prenatal nutrition on kidney development and its link to chronic kidney disease and hypertension in adulthood. By identifying specific gene expression changes and pathways affected by LPD, this research offers potential targets for therapeutic intervention and underscores the importance of adequate maternal nutrition during pregnancy. Moreover, this study addresses previously unanswered questions regarding the specific cellular and molecular disruptions caused by LPD, and how reduced protein intake during critical developmental windows leads to deficiencies in nephron endowment. These insights contribute to a more comprehensive understanding of the developmental origins of health and disease, informing both clinical approaches and public health strategies to mitigate the impact of prenatal nutritional deficits on long-term health outcomes.

## Materials and Methods

### IACUC statement

All animal experiments in this study were assessed and approved by the Monash University Animal Ethics Committee (MARP/18204) and the Tulane University Institutional Animal Care and Use Committee (1558). Animal work was conducted under applicable laws governing the care and use of animals for scientific purposes.

### Animals and diets

CD1 mice were acquired from Charles River Laboratories and C67BL6/J mice were sourced from Jax Laboratories via Monash University. Six-week-old females were fed either Normal (20-18%) protein diet (ENVIGO, cat# TD.91352, Specialty Feeds, cat# AIN-93G respectively) or Low (6-8%) protein diet (Specialty Feeds, cat# SF01-026, ENVIGO, cat# TD.90016) for 2 weeks. International transport restrictions between the El-Dahr and Smyth groups located in the USA and Australia respectively determined the differences between the diets. See Supplemental Dataset 1 for diet composition. While there are differences and similarities between the diets and animal strains, the kidney phenotype presented in the present study is the same in the LPD exposed offspring.

### Animal collection

Time mated pregnant females were euthanized at approximately E14.5, E19.5 (day of vaginal plug being E0.5), and pups euthanized at P0 and P2. Mouse embryos and pups were staged according to limb staging ^46^, dissected and the kidneys were removed and either fixed in 4% formalin and put in ice-cold PBS, or liquid nitrogen (depending on experimental requirements).

### Niche assessment

Dissected kidneys at E14.5, P0 and P2 were stained with antibodies to NPCs (SIX2, Proteintech cat#: 115621AP, 1:600 dilution), ureteric tree (TROP2, R&D Systems, cat#: AF1122, 1:20 dilution), proliferating nuclei (PHH3, Abcam, cat#: AB10543, 1:300 dilution) and nuclei (DAPI, Sigma, cat#: D9542, 1mg/mL dilution) with appropriate Alexa Fluor secondary antibodies (Life Technologies) as previously described ^47^. The stained kidneys were cleared using BABB (Benzyl alcohol/Benzyl benzoate, 1:2, Sigma, cat numbers 108006 and B6630, consecutively) and imaged using a Leica SP8 confocal microscopy at 20X and quantified using Imaris (Oxford Instruments). In order to accurately sample the cellular state across the kidney, morphological assessments were made for per-tip and per-cap (i.e. per niche) with a minimum 6 tip/cap combinations per imaged field, with usually 6 fields per kidney covering both sides, as well as at least one pole (less fields were available at E14.5 where the kidneys are smaller). A grand total of 78 niches at E14.5 (from 8 kidneys), 564 at P0 (from 20 kidneys), and 289 at P2 (from 14 kidneys) were assessed. Cell numbers, proliferating cell numbers, cap-mesenchyme, and tip volumes were all measured for each niche, as well as an assessment of associated nephric structures. Nephric structure assessment was qualitatively performed by observation of nuclei in Z slices near ureteric tips, and defining aggregates, luminal balls of cells, or elongated tubular luminal structures of different lengths and morphologies. This enabled the distinction between nephron progenitor cells, pretubular aggregates, renal vesicles, comma/S-shaped bodies, and nephrons.

### Histological staining

Dissected kidneys at E14.5, P0 were fixed and histologically sectioned and were stained with antibodies to detect Wnt4 (anti-Wnt4, R&D systems, cat# AF475, dilution 1:100), Wnt9b (anti-Wnt9b, R&D Systems, cat# AF3669, 1:100 dilution), Six2 (as above but at 1:400 dilution) in different combinations. The primary markers were detected with secondary antibodies using HRP (Jackson Immuno Research, cat# 705-036-174, 1:100 dilution), Alexa Fluor 555 (Thermo Fisher, cat# A-31572, 1:400 dilution), fluorescent tagged Dolichos Biflorus Agglutinin (DBA) (Vector Labs, cat# FL1031, 1:100 dilution), and nuclei with Hoechst-33342 (Thermo Fisher, cat# 62249, dilution 1:10,000 dilution).

### Optical Projection tomography, branching, and niche number analysis

TROP2 antibody stained E14.5 kidneys were mounted in agarose, cleared, and imaged in a Bioptonics OPT3001 instrument using 800 projections at 1024x1024 pixel resolution as previously described ^47^. Tomography projections were reconstructed using nRecon (Bruker) and Tree Surveyor was used to map the ureteric tree as previously described ^21, 47^ which enabled the determination of branch lengths, surface area, kidney volume, tip number and tip to tip distances. Branch patterning was quantified from Tree Surveyor data using python scripts as previously described ^22^. To count tip/niche number at P0, whole kidneys were stained with SIX2 antibody (1:600), and an appropriate Alexa-fluor secondary antibody (Life Technologies), cleared, and imaged using optical projection tomography as above. SIX2 positively stained clusters/niches were classified by their characteristic (‘horseshoe’) shape and manually counted in Imaris (Oxford Instruments). Each cluster represents a single niche, and by extension a single ureteric/collecting duct tip ^32^.

### Statistics

In order to statistically test the differences between diets where repeated measurements are taken per-kidney (e.g., proliferation or associated nephric structures per niche from confocal data, body mass over time, or branching measurements from OPT/Tree Surveyor), linear mixed-effects models were applied to the data within the R statistical environment using the stats or lme4 packages. The use of two factor ANOVA enabled the partitioning of inter-kidney variance within the treatment group. To quantify the cumulative probability of differential association of cells associated across pseudotime, we used a Two-sample KS (Kolmogorov-Smirnov) test. Two sample tests were performed using an unpaired two-tailed T-test.

### Nephron counting

NPD and LPD offspring were collected at P0 to assess the impact of gestational LPD feeding on nephron number. Pups were weighed, sexed and kidneys dissected in PBS and fixed in methacarn (60% absolute methanol, 30% chloroform, and 10% glacial acetic acid). Following processing to paraffin, the kidneys were exhaustively sectioned at 4 µm. Ten evenly spaced section pairs were systematically sampled and stained with lectin peanut agglutinin (PNA; Sigma-Aldrich, Castle Hill, NSW, Australian; L3165) to identify the plasma membranes of podocytes. Following counterstaining with hematoxylin, section pairs were projected using a light microscope and all PNA+ structures (representing S-shaped body stage to capillary loop stage glomeruli) were counted using the physical dissector/fractionator counting principle ^48^. The same approach was applied at P21, except the section thickness was increased to 5 µm.

### Single-cell and nuclear RNA-sequencing and data analysis

Two groups of kidneys five E14.5 kidneys, from 2 different females, per diet were pooled together and processed to obtain single-cell suspension as previously described ^49^. For single nuclear sequencing at postnatal day 0, pairs of kidneys were snap frozen in liquid nitrogen immediately after dissection. Single nuclear approach was specifically chosen for the postnatal kidneys because they require significantly more physical and enzymatic intervention for cellular dissociation than E14.5 kidneys, and this treatment may significantly impact cellular and sequencing quality. Frozen nuclei were thawed in nuclei lysis buffer (Sigma) in a 15mL tissue grinder (Kimble), ground, and isolated by centrifugation and washing before being purified by flow cytometry, as per ^50^. Approximately 10k cells or nuclei (per sample) were emulsified with Single Cell 3’ V3 beads (10X Genomics) and libraries generated on a 10X Chromium instrument. Libraries were amplified and sequenced using an MGITech MGISEQ200RS with a target of 50k reads per cell. Raw sequencing data was processed using Cell Ranger (v5.0, 10X Genomics) and transcripts mapped to a reference genome based on mm10 (v3) with pre-mRNA (nuclear) optimizations. Single cell preparation and sequencing of E14.5 samples was performed using published embryonic kidney dissociation methods ^49^ and using protocols from 10X Genomics. Single-cell samples were sequenced with an Illumina NextSeq 2000 mRNA (cellular), and processed as above, but with mapped to mm10 (v3) genome mRNA. All sequencing analysis was performed in R with the aid of the Tidyverse packages ^51^. Seurat (v3.2) ^52^ was the major package used to import and analyze the cell/nuclear sequencing data. Cells with no counts or less than 300 genes detected, or more than 15% of mitochondrial genes were removed. Supervised clustering was aided by Clustree ^53^ and cluster identity was determined using 4000 highly variable genes, and the first 20 principal components. We obtained lists of DE genes for each cluster using the Seurat FindAllMarkers function and upregulated genes in each cluster were used to classify the different cell types. To test differential gene expression between diets, Seurat objects were converted to single-cell experiment objects and the limma package ^54^ was used for testing. To further understand gene biological functions and associated pathways, we performed the gene ontology (GO) and KEGG pathway enrichment analysis with Goanna (from limma), and additional unsupervised and supervised ontology analysis with STRING and DAVID ^55^, and GOfuncR ^56^ for extracting gene annotations from ontologies. Pseudotime analysis was performed using Slingshot ^57^, and differential gene expression analysis across pseudotime was performed using TradeSeq^58^. The sequencing datasets used in this work are available on GEO accession ID GSE240044.

## Acknowledgments

We would like to acknowledge the many contributors to this work including technical assistance from Grace Jackel and Guizhi Sun at Monash University. We also would like to thank James LeFevre (University of Queensland) and Alexander Combes (Monash University) for bioinformatic and data processing discussions. We thank staff and managers of the facilities used for this work, including Monash Animal Research Platform, Monash Micro Imaging, Monash FlowCore, Monash Micromon Genomics, Monash Histology Platform, the Monash Bioinformatics core, the Tulane Center for Translational Research in Infection & Inflammation NextGen Sequencing Core. This work was funded by Tulane School of Medicine Startup Grant to Dr. Tortelote, NIDDK R01 grant (5R01DK118231) to S. El-Dahr, and NHMRC project grant APP1080746 to Profs. Smyth, Bertram and Belwitt.

