## Supplementary figures and images for "The Impact of Low Protein Diet on the Molecular and Cellular Development of the Fetal Kidney"

### Supplementary Figure 1

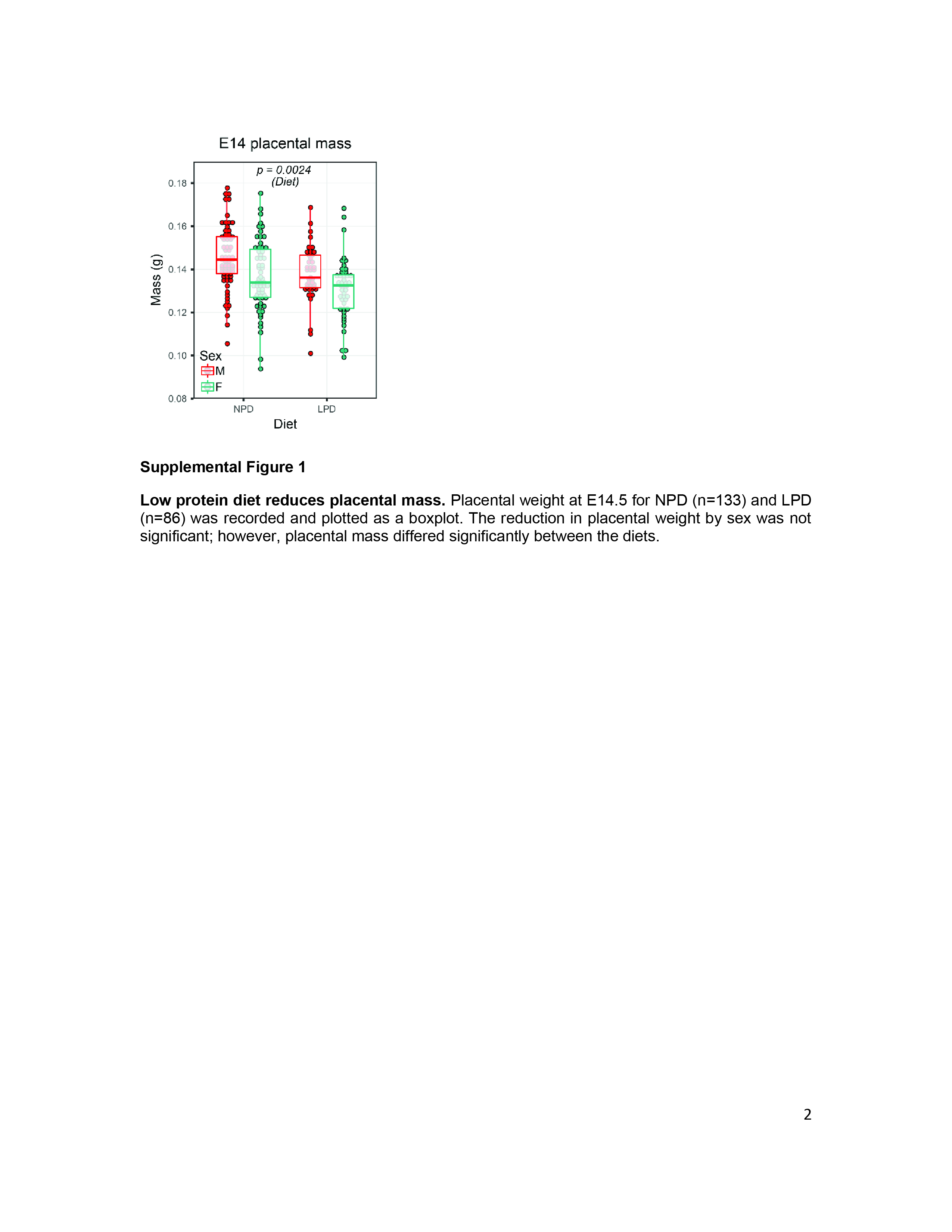

### Supplementary Figure 2

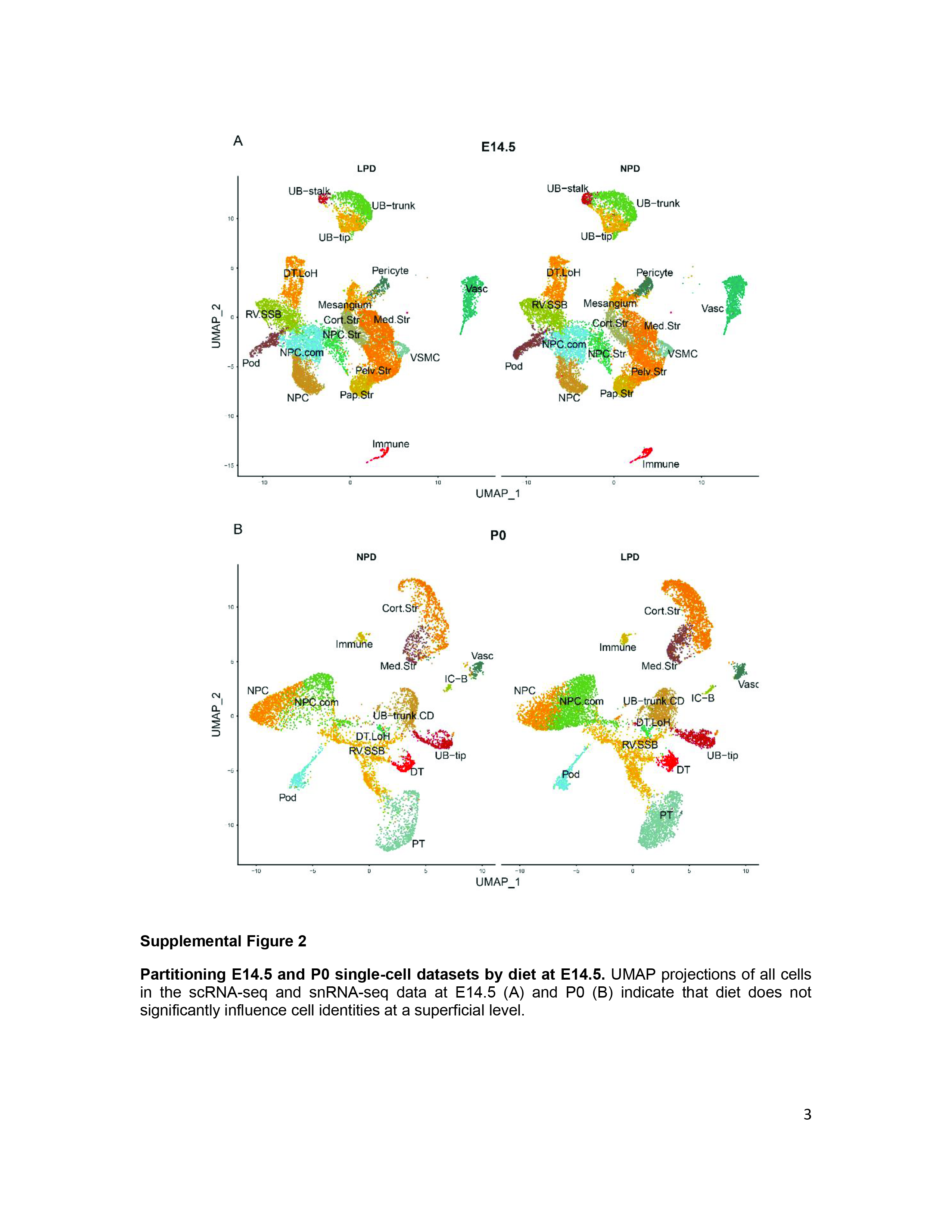

### Supplementary Figure 3

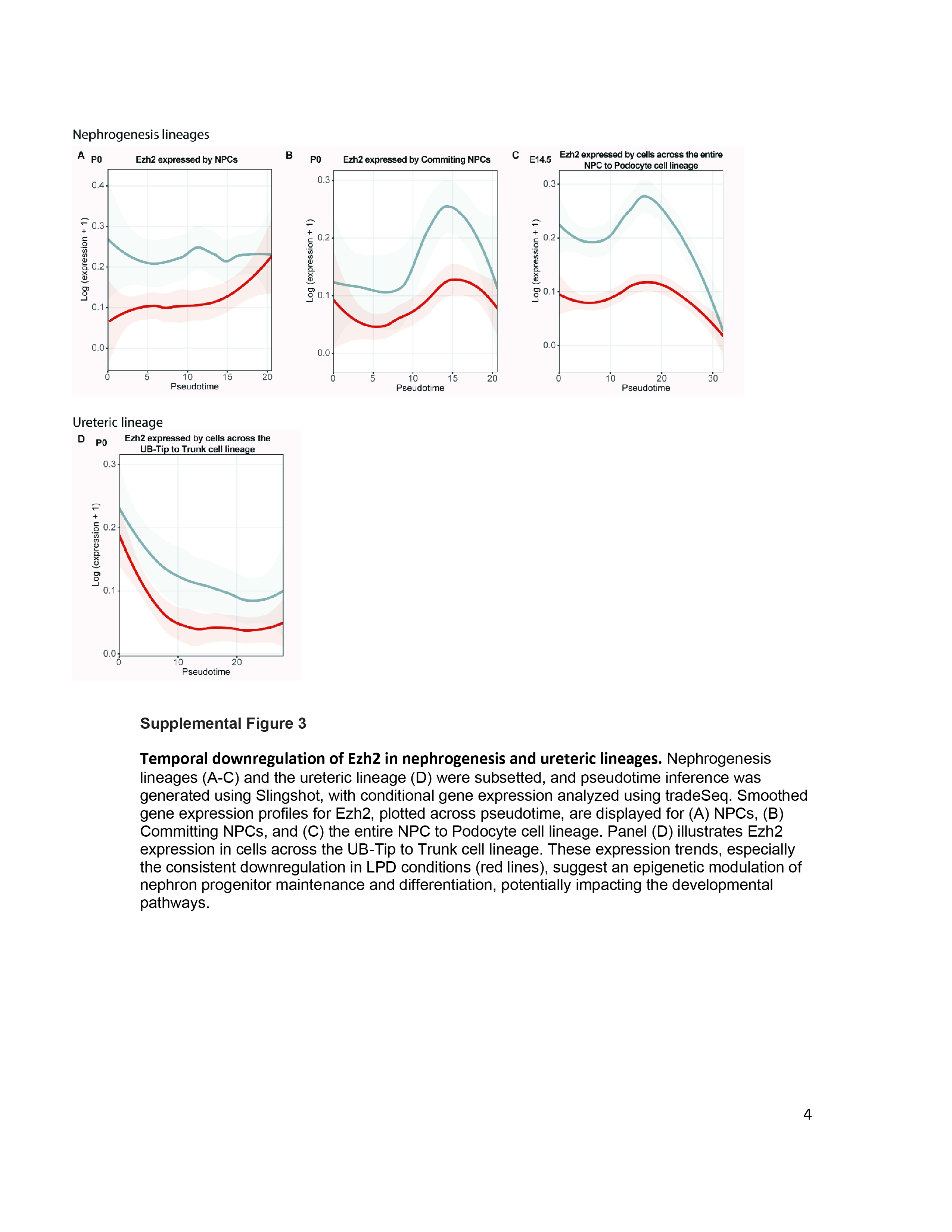

### Supplementary Figure 4

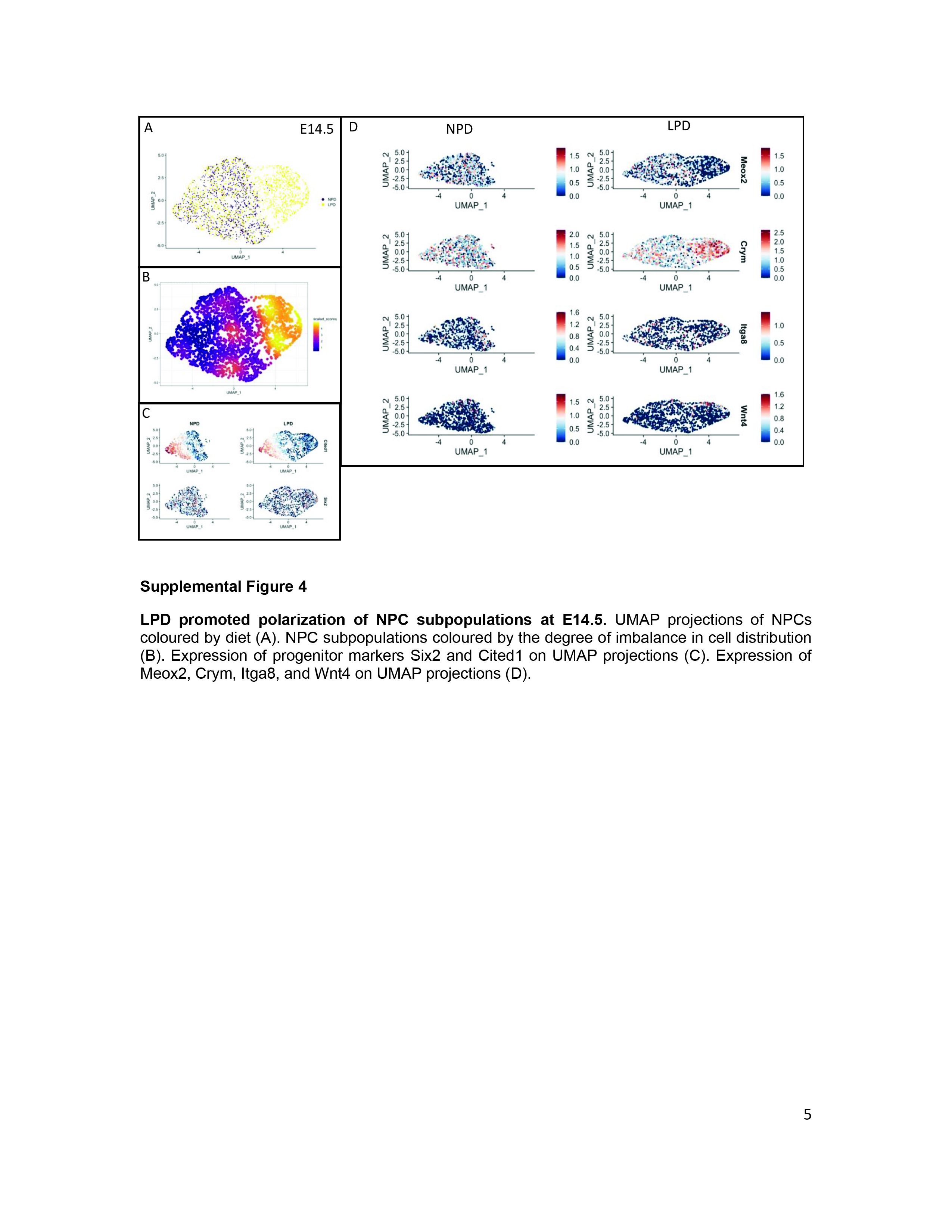

### Supplementary Figure 5

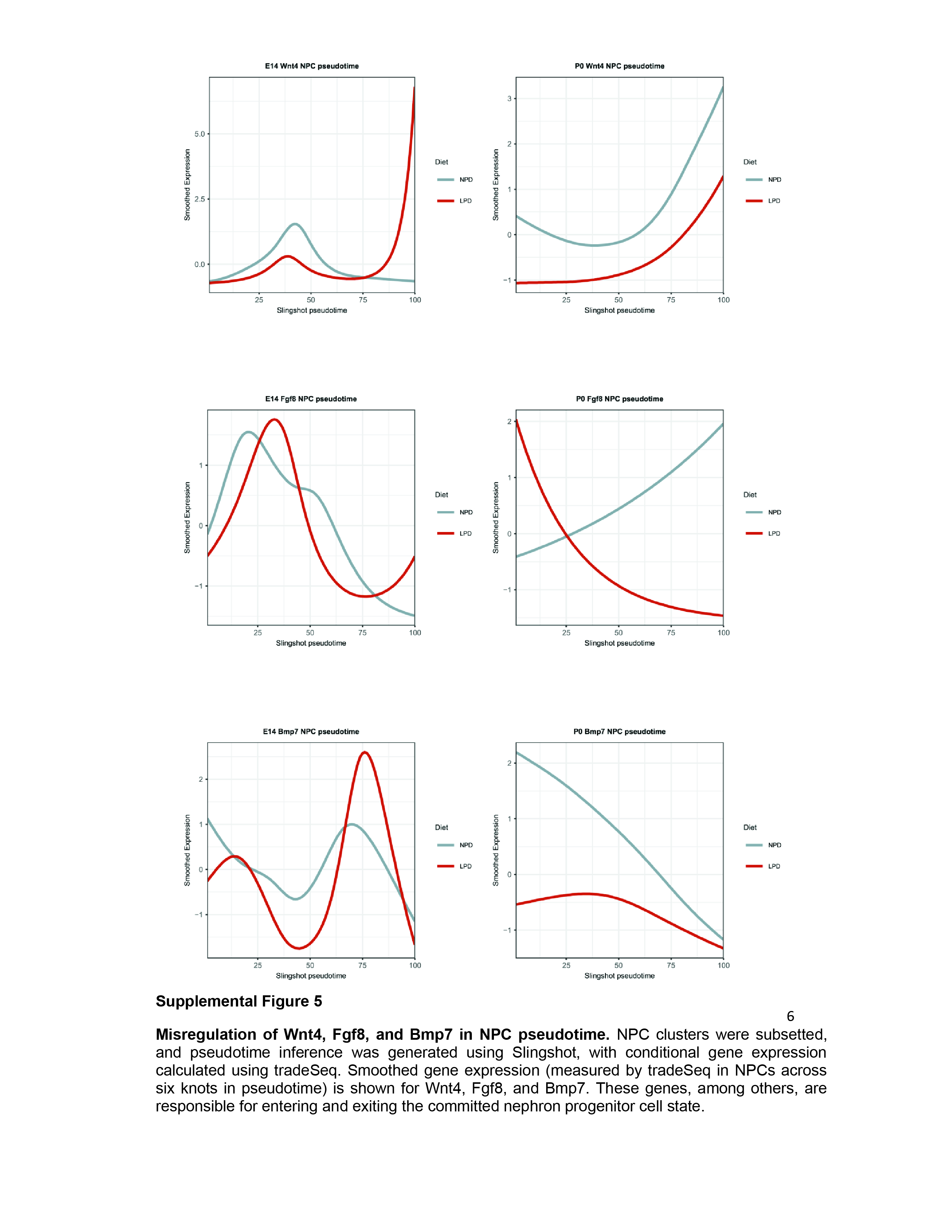

### Supplementary Figure 6

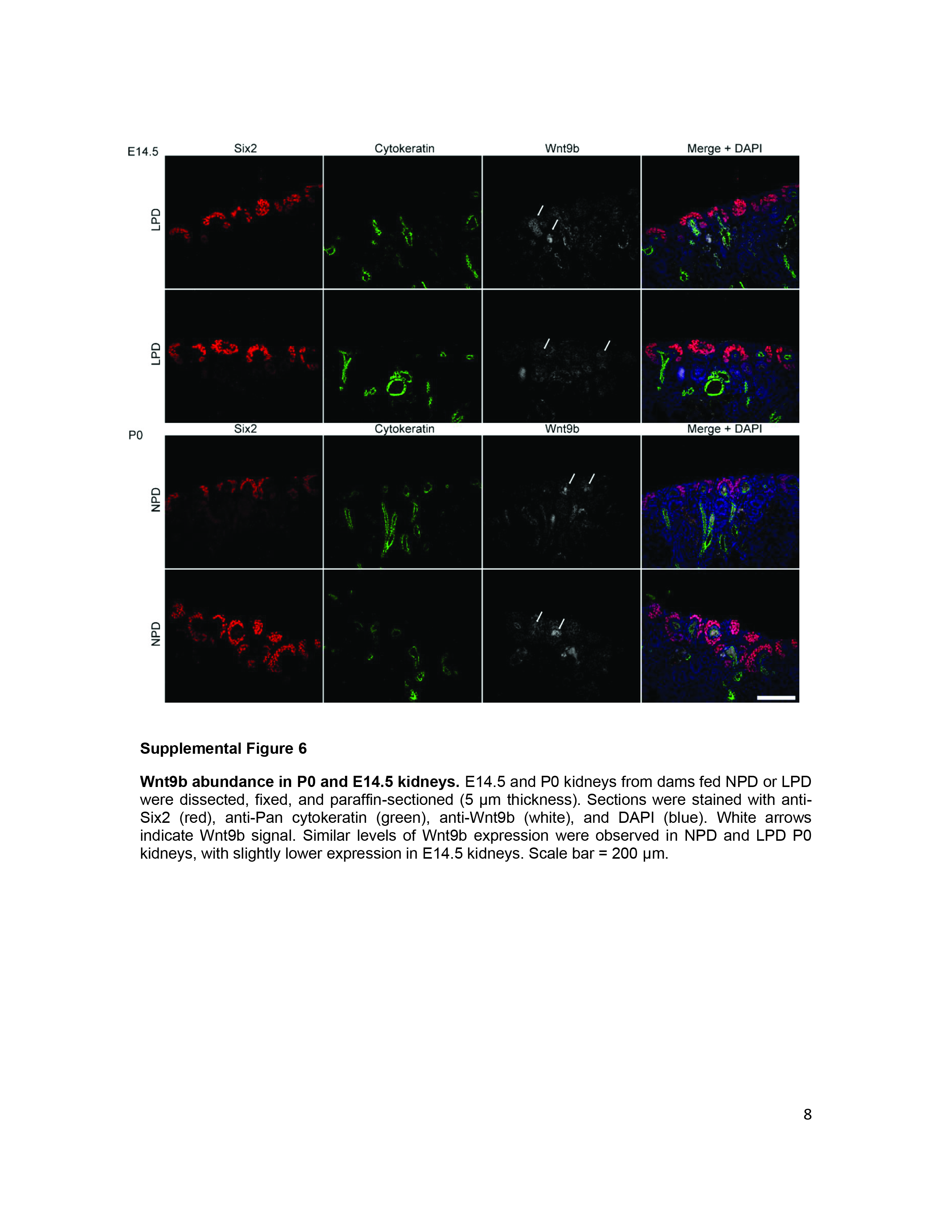
