## Supplementary Figure Legends for "The Impact of Low Protein Diet on the Molecular and Cellular Development of the Fetal Kidney"

**Supplemental Figure 1**

**Low protein diet reduces placental mass.** Placental weight at E14.5 for NPD (n=133) and LPD (n=86) was recorded and plotted as a boxplot. The reduction in placental weight by sex was not significant; however, placental mass differed significantly between the diets.

**Supplemental Figure 2**

**Partitioning E14.5 and P0 single-cell datasets by diet at E14.5.** UMAP projections of all cells in the scRNA-seq and snRNA-seq data at E14.5 (A) and P0 (B) indicate that diet does not significantly influence cell identities at a superficial level.

**Supplemental Figure 3**

**Temporal downregulation of Ezh2 in nephrogenesis and ureteric lineages.** Nephrogenesis lineages (A-C) and the ureteric lineage (D) were subsetted, and pseudotime inference was generated using Slingshot, with conditional gene expression analyzed using tradeSeq. Smoothed gene expression profiles for Ezh2, plotted across pseudotime, are displayed for (A) NPCs, (B) Committing NPCs, and (C) the entire NPC to Podocyte cell lineage. Panel (D) illustrates Ezh2 expression in cells across the UB-Tip to Trunk cell lineage. These expression trends, especially the consistent downregulation in LPD conditions (red lines), suggest an epigenetic modulation of nephron progenitor maintenance and differentiation, potentially impacting the developmental pathways.

**Supplemental Figure 4**

**LPD promoted polarization of NPC subpopulations at E14.5.** UMAP projections of NPCs coloured by diet (A). NPC subpopulations coloured by the degree of imbalance in cell distribution (B). Expression of progenitor markers Six2 and Cited1 on UMAP projections (C). Expression of Meox2, Crym, Itga8, and Wnt4 on UMAP projections (D).

**Supplemental Figure 5**

**Misregulation of Wnt4, Fgf8, and Bmp7 in NPC pseudotime.** NPC clusters were subsetted, and pseudotime inference was generated using Slingshot, with conditional gene expression calculated using tradeSeq. Smoothed gene expression (measured by tradeSeq in NPCs across six knots in pseudotime) is shown for Wnt4, Fgf8, and Bmp7. These genes, among others, are responsible for entering and exiting the committed nephron progenitor cell state.

**Supplemental Figure 6**

**Wnt9b abundance in P0 and E14.5 kidneys.** E14.5 and P0 kidneys from dams fed NPD or LPD were dissected, fixed, and paraffin-sectioned (5 μm thickness). Sections were stained with anti-Six2 (red), anti-Pan cytokeratin (green), anti-Wnt9b (white), and DAPI (blue). White arrows indicate Wnt9b signal. Similar levels of Wnt9b expression were observed in NPD and LPD P0 kidneys, with slightly lower expression in E14.5 kidneys. Scale bar = 200 μm.
